# A hypomorphic allele of telomerase reverse transcriptase uncovers the minimal functional length of telomeres in Arabidopsis

**DOI:** 10.1101/520163

**Authors:** J. Matthew Watson, Johanna Trieb, Martina Troestl, Kyle Renfrew, Terezie Mandakova, Dorothy E. Shippen, Karel Riha

## Abstract

Despite the essential requirement of telomeric DNA for genome stability, the length of telomere tracts between species differs by up to four orders of magnitude, raising the question of the minimal length of telomeric DNA necessary for proper function. Here we address this question using a hypomorphic allele of the telomerase catalytic subunit, TERT. We show that although this construct partially restored telomerase activity to a *tert* mutant, telomeres continued to shorten over several generations, ultimately stabilizing at a bimodal size distribution. Telomeres on two chromosome arms were maintained at a length of 1kb, while the remaining telomeres were maintained at 400 bp. The longest telomeres identified in this background were also significantly longer in wild type populations, suggesting cis-acting elements on these arms either promote telomerase processivity or recruitment. Genetically disrupting telomerase processivity in this background resulted in immediate lethality. Thus, telomeres of 400 bp are both necessary and sufficient for Arabidopsis viability. As this length is the estimated minimal length for t-loop formation, our data suggest that telomeres long enough to form a t-loop constitute the minimal functional length.

## Introduction

The ends of most eukaryotic chromosomes are capped by telomeres that serve two primary functions: they distinguish the natural chromosome end from a DNA double-strand break (DSB) and provide a mechanism for overcoming the end-replication problem (Fouquerel, Parikh et al., 2016, Stewart, Chaiken et al., 2012). Telomeres are composed primarily of repetitive telomeric DNA and telomere specific DNA binding proteins (de Lange, 2010). The telomeric repeat is a short G-rich sequence, TTAGGG in vertebrates and TTTAGGG in plants (Fulcher, Derboven et al., 2014). Telomeric DNA cannot be fully replicated by conventional DNA polymerases, an issue known as the end-replication problem. During lagging strand synthesis, removal of the final RNA primer results in the formation of a gap in the DNA that cannot be replaced, leading to the formation of a 3’ single-stranded DNA overhang (G-overhang) and a small loss of terminal sequences (Soudet, Jolivet et al., 2014). The end-replication problem can be counteracted by the enzyme telomerase, consisting minimally of the catalytic subunit TERT (telomerase reverse-transcriptase) and an internal RNA template, TER (telomerase RNA) (Greider & Blackburn, 1989). Telomerase can anneal to the G-overhang and, using its internal template, processively extend the extreme 3’ end (Tomita, 2018).

Telomeric repeats serve as a platform for sequence-specific telomere binding proteins that can associate either with double- or single-stranded DNA. In mammals, double-strand telomere binding proteins TRF1 and TRF2 anchor a multi-subunit protein complex termed shelterin to telomeres (de Lange, 2010). Shelterin is necessary for preventing cells from recognizing telomeres as DSBs (Sfeir & de Lange, 2012), but precisely how this is accomplished is unknown. One model predicts that chromosome ends are protected through the formation of t-loops. T-loops are higher order structures in which the G-overhang invades duplex telomeric DNA, forming a looped molecule that hides the chromosome 3’ terminal extension (Doksani, Wu et al., 2013, Griffith, Comeau et al., 1999). Although first described in mammals, T-loops have also been identified in other taxa including plants (Cesare, Quinney et al., 2003, Munoz-Jordan, Cross et al., 2001, Raices, Verdun et al., 2008, Tomaska, Makhov et al., 2002). T-loops likely do not form in organisms that possess relatively short telomeres, such as *Saccharomyces cerevisiae* or in the vegetative nuclei of ciliates (Luke-Glaser, Poschke et al., 2012, Tomaska, Willcox et al., 2004). In these species, chromosome end protection appears to largely depend on protein complexes associated with single-stranded telomeric DNA (Greetham, Skordalakes et al., 2015).

In the absence of telomerase, telomeric DNA gradually shortens, eventually leading to telomere dysfunction. Thus, the ability of telomeres to protect chromosome ends and allow for cellular viability is a function of their length. The molecular details of this relationship are unclear. Telomeres are maintained at different lengths in different organisms, ranging from 300 bp in yeasts to as long as 150 kb in tobacco and the telomere length can also be highly variable within a species (Fajkus, Kovarik et al., 1995, Fulcher, Teubenbacher et al., 2015, Liti, Haricharan et al., 2009, Shakirov & Shippen, 2004). Most organisms keep telomeres at lengths longer than necessary for function to provide a protective buffer, as defects in telomere maintenance are often only apparent after multiple generations (Blasco, Lee et al., 1997, Lundblad & Szostak, 1989, Meier, Clejan et al., 2006, Nakamura, Morin et al., 1997, Riha, McKnight et al., 2001). Several attempts have been made to determine the minimal length of telomeric DNA necessary for telomere function (Capper, Britt-Compton et al., 2007, Heacock, Spangler et al., 2004, Xu & Blackburn, 2007). These approaches have relied on the identification of the smallest telomere within a population undergoing telomere crisis due to ongoing telomere shortening. The minimal functional lengths inferred from these experiments vary from roughly 100 bp in yeast (Forstemann, Hoss et al., 2000, Teixeira, Arneric et al., 2004), 42-78 bp in human cell lines (Capper et al., 2007, Xu & Blackburn, 2007), and 360 bp in Arabidopsis (Heacock et al., 2004). However, it has not been possible to determine whether the very short telomeres identified in these studies are actually functional, since they form only a small portion of total telomeres and may represent dysfunctional and nucleolytically processed intermediates. Because variations in minimal telomere length may belie fundamental differences in telomere capping, determining the amount of telomeric DNA necessary for telomere function can provide insight into mechanisms governing chromosome-end protection.

Here we report the creation of Arabidopsis strains that maintain their telomeres at the minimal functional threshold by expressing a hypomorphic allele of TERT. While plants with these shortened telomeres exhibit chromosome end-to-end fusions and ongoing genome instability, they are capable of indefinite survival as long as they are not subjected to further telomere erosion. Thus, we propose that the final telomere length reached in this population, roughly 400 bp, represents the minimal length necessary for proper telomere function in Arabidopsis.

## Results

Following the discovery and analysis of telomerase mutants in Arabidopsis, we attempted to study the fate of shortened telomeres upon reintroduction of telomerase activity. To this end, we transformed 4^th^ generation *tert* mutants (G4) with constructs expressing the *TERT* cDNA under the control of either the cauliflower mosaic virus 35S or actin (ACT2) promoter (Fig. S1). While both constructs restored telomerase activity, TRAP assays revealed that enzyme activity was significantly reduced compared to wild type plants. Plants bearing the *ACT2:TERT* construct showed slightly higher levels of telomerase than the *35S:TERT* lines (Fig. 1A).

**Fig 1.**
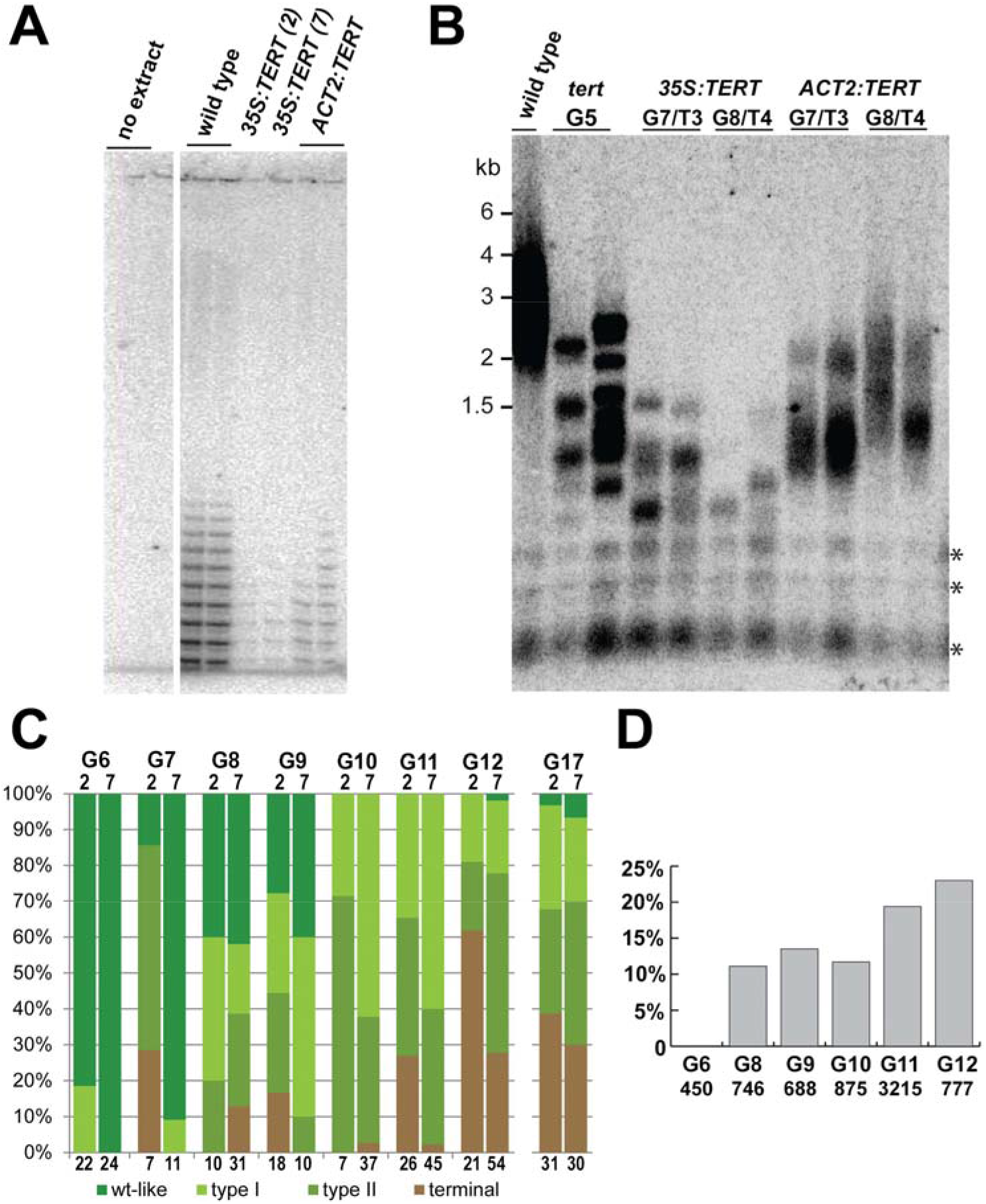
Phenotypic analysis of the *35S:TERT* allele. (*A*) TRAP assay of wild type seedlings and seedlings transformed with either *35S:TERT* or *ACT2:TERT* constructs. Both constructs show reduced telomerase activity relative to wild type. (*B*) TRF analysis of successive generations of *35S:TERT* or *ACT2:TERT* plants. Wild type and *tert* mutants are shown as controls; the asterisk marks interstitial telomeric DNA. Rescue constructs were transformed into G4 *tert* mutants. The “T” designation indicates the number of generations post transformation (Fig S1). (*C*) Phenotypic analysis of successive generations of plants containing the *35S:TERT* construct. Two independent lines were analyzed (line 2 and 7) with G6/T2, G7/T3 etc. The total number of plants analyzed in each population is indicated below the bars. (*D*) Percentage of anaphases containing chromatin bridges over successive generations of plants containing the *35S:TERT* construct. Total number of anaphases examined is indicated. Data represents pools from both lines.

We followed two independent transformant lines for both constructs through multiple generations (Fig. S1) and measured telomere length by terminal restriction fragment length analysis (TRF) in G7/T3 and G8/T4. Telomeres in the two lines expressing *TERT* driven by the actin promoter showed a gradual lengthening of telomeres (Fig. 1B), consistent with restoration of telomerase function in telomere synthesis. In contrast, both lines in which *TERT* expression was driven by the 35S promoter exhibited continual shortening of telomeres. The telomeric signal was under 1.5 kb in G7/T3 in *35S:TERT* plants and telomeres further shortened in G8/T4 (Fig. 1B). Telomeres were also substantially shorter than telomeres of parental G5 *tert* plants. Additionally, telomeres in the *35S:TERT* lines displayed a discrete TRF banding pattern, a feature typical for plants with inactive telomerase. These data indicate that despite detectable enzyme activity, telomerase in the *35S:TERT* lines is incapable of counteracting the end-replication problem and telomeres continue to shorten.

To follow the fate of lines expressing the hypomorphic TERT allele, we continued to propagate the two *35S:TERT* lines, named 2 and 7, for multiple generations. Unexpectedly, despite the progressive loss of telomeric DNA observed in the initial TRFs (Fig. 1), we were able to self-propagate both line 2 and line 7 through generation G18/T14, when the experiment was discontinued. This is ten generations longer than *tert* null mutants, which arrest growth in a terminal generation of G8 - G10 (Riha et al., 2001). Notably, out-segregation of the *35S:TERT* construct in G7/T3 resulted in termination of the population in the next generation (Fig. S1), confirming that the *35S:TERT* construct was responsible for survival of complemented *tert* mutants. Despite the ability of the *35S:TERT* lines to survive for multiple generations, they began to display phenotypes characteristic of genomic instability within two generations after transformation (G7/T3, Fig. 1). These phenotypes including wrinkled leaves, fasciated stems, altered phyllotaxy, and reduced seed set. Plants with these phenotypes have previously been categorized based on increasing severity into Type I, Type II, and Terminal phenotypes (Riha et al., 2001), with Type I plants showing mild defects in leaf morphology and phyllotaxy, Type II plants exhibiting severe developmental defects and reduced fertility, and dwarfed terminal plants that do not produce any seeds. We scored populations of plants from both *35S:TERT* lines for the frequency of these phenotypic categories over multiple generations (Fig. 1C). The severity of the phenotypes increased with each successive generation until G12/T8, indicative of increasing telomere dysfunction. In accordance with the increasing severity of the phenotypes, we observed a gradual increase in the frequency of anaphase bridges through G12/T8 (Fig. 1D), suggesting that the worsening phenotypes are due to increased genomic instability caused by telomere erosion. Notably, the fraction of plants within the respective phenotypic categories became stabilized in subsequent generations after G12/T8, with approximately one third of plants exhibiting wild type or Type I phenotypes (Fig. 1C and Fig. S2). This observation suggests that surviving populations exhibit ongoing genome instability and partial telomere dysfunction.

Several reports have indicated that telomerase has biological functions beyond its primary role of extending telomeres (Blasco, 2002, Li & Tergaonkar, 2014). To test whether the extended lifespan of the *35S:TERT* lines is due to a non-canonical function of telomerase, we generated a *TERT* cDNA construct containing a point mutation (D860N) in a conserved catalytic residue necessary for telomerase function (Nakayama, Tahara et al., 1998). This construct was transformed into G4 *tert* plants, and two independent transformants were analyzed for telomerase activity and long-term survival. As expected, we were unable to detect telomerase activity from the *35S:TERT(D860N)* lines (Fig. S3). The plants carrying the catalytically dead *TERT* construct survived for only two generations after transformation, essentially phenocopying the parental *tert* mutants (Fig S1, Fig. 2A). We conclude that the continued survival of *35S:TERT* plants is due to telomerase activity, and not due to a non-canonical function of the TERT protein.

**Fig 2.**
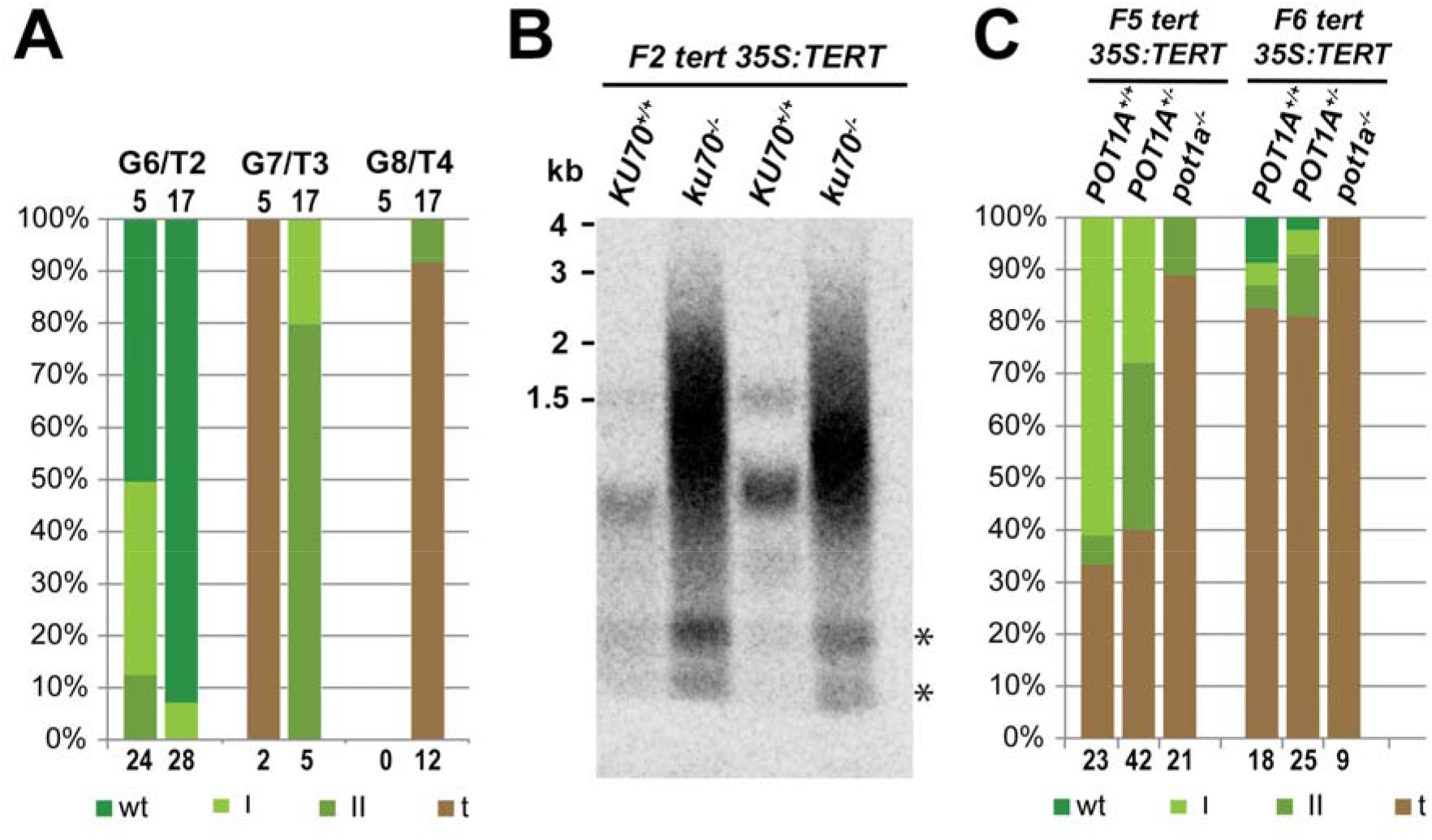
Genetic analysis of the *35S:TERT* allele. (*A*) Phenotypic analysis of successive generations of *35S:TERT(D680N)* transformants. In the absence of telomerase catalytic activity, two independent transformant lines (5 and 17) show rapid loss of viability. (*B*) TRF analysis of *35S:TERT* containing siblings segregating for *ku70* (see Fig. S4 for details of the cross). Loss of *KU70* leads to immediate lengthening of telomeres. Asterisks mark interstitial telomeric DNA. (*C*) Phenotypic analysis of *35S:TERT* plants segregating for *POT1A*. See Fig. S5 for details for the cross. Loss of POT1A increases severity of the phenotypes in *35S:TERT* background.

To determine whether the *35S:TERT* construct acts through the canonical telomere maintenance pathway, we examined its genetic interaction with other factors known to be involved in regulating telomerase activity at telomeres in Arabidopsis. We first asked whether the *35S:TERT* allele could extend telomeres in the absence of KU70, a negative regulator of telomerase whose inactivation results in telomerase-dependent telomere lengthening (Riha & Shippen, 2003, Riha, Watson et al., 2002). We generated *ku70 tert 35:TERT* plants by crossing (Fig. 2A and S4). Loss of KU70 in the *35S:TERT* background led to an immediate and obvious lengthening of telomeres, demonstrating that despite its reduced activity (Fig. 1A), the *35S:TERT* telomerase is capable of extending telomeres. Next, we asked whether survival of the *35S:TERT* lines depends on POT1A, an OB-fold containing protein that directly interacts with the telomerase RNP and is required for telomerase repeat addition processivity (Renfrew, Song et al., 2014, Surovtseva, Shakirov et al., 2007). Plants deficient for POT1A exhibit an ever-shorter telomere phenotype, and this phenotype is epistatic with TERT (Surovtseva et al., 2007). We crossed third generation *tert pot1a* double mutants to T2 *35S:TERT* plants from line 2 (Fig. S5), and propagated the progeny for an additional three generations as *POT1A^+/−^*, allowing the longer telomeres originating from the *tert pot1a* parent to shorten. We then outsegregated *pot1a* mutants in F5 and F6 and scored the population for growth phenotypes (Fig. S5 and 2C). Population of *POT1A^−/−^* mutants in both generations showed more severe phenotypes than either the *POT1A^+/+^* or *POT1A^+/−^* plants, and all *pot1a^−/−^* plants in the F6 population displayed terminal phenotype. These results demonstrate that the survival of *35S:TERT* lines requires POT1A.

Taken together, these genetic experiments suggest that the long-term survival of the *35S:TERT* lines depends on the ability of telomerase to add telomeric repeats to the telomeres. We hypothesized that the *35S:TERT* construct leads to the formation of a hypomorphic telomerase enzyme that is unable to rescue the shortening of telomeres in *tert* mutants, but supports continuous growth in a significant fraction of the population by maintaining telomeres once they reach minimal length necessary for telomere protection. To test this prediction, we analyzed telomeres in *35S:TERT* lines by PETRA, a PCR based method that permits precise analysis of telomere length at individual chromosome arms (Heacock et al., 2004). We measured the length of telomeres at seven chromosome arms in four to five sibling plants from each line, every two generations, between G6 /T2 and G14/T10 (Fig. 3A). The plants selected for analysis were randomly sampled from the pool of plants displaying Type I and Type II phenotypes. At the beginning of the experiment in G6/T2, telomeres ranged in size from a low of 285 bp to a high of 1797 bp with an average length of 951 ± 350 bp. In line 2, telomeres that began with a length above 1 kb shortened at an average rate of 243 ± 102 bp per generation, and this rate continued through G10/T6 in the longest telomeres (1R, 2R, and 5L). We observed a similar trend in line 7, where telomeres longer than 1kb (1L, 2R, and 5L) shortened with an average rate of 198 ± 80 bp per generation. A similar rate of telomere shortening has been reported for *tert* mutants (Riha et al., 2001, Watson & Shippen, 2007), suggesting that the hypomorphic telomerase is not extending these telomeres. In contrast to the longest telomeres, the shortest telomeres in both lines remained essentially unchanged throughout the eight generations of the experiment (Fig. 3A, 1L and 5R in line 2, 5R in line 7), indicating that they are sustained at these lengths by the hypomorphic telomerase. Additionally, while the longest telomeres shortened quite rapidly between G6/T2 and G10/T6, the rate of shortening became negligible from G10/T6 onwards, and telomere length was stable for the remaining four generations of the experiment.

**Fig 3.**
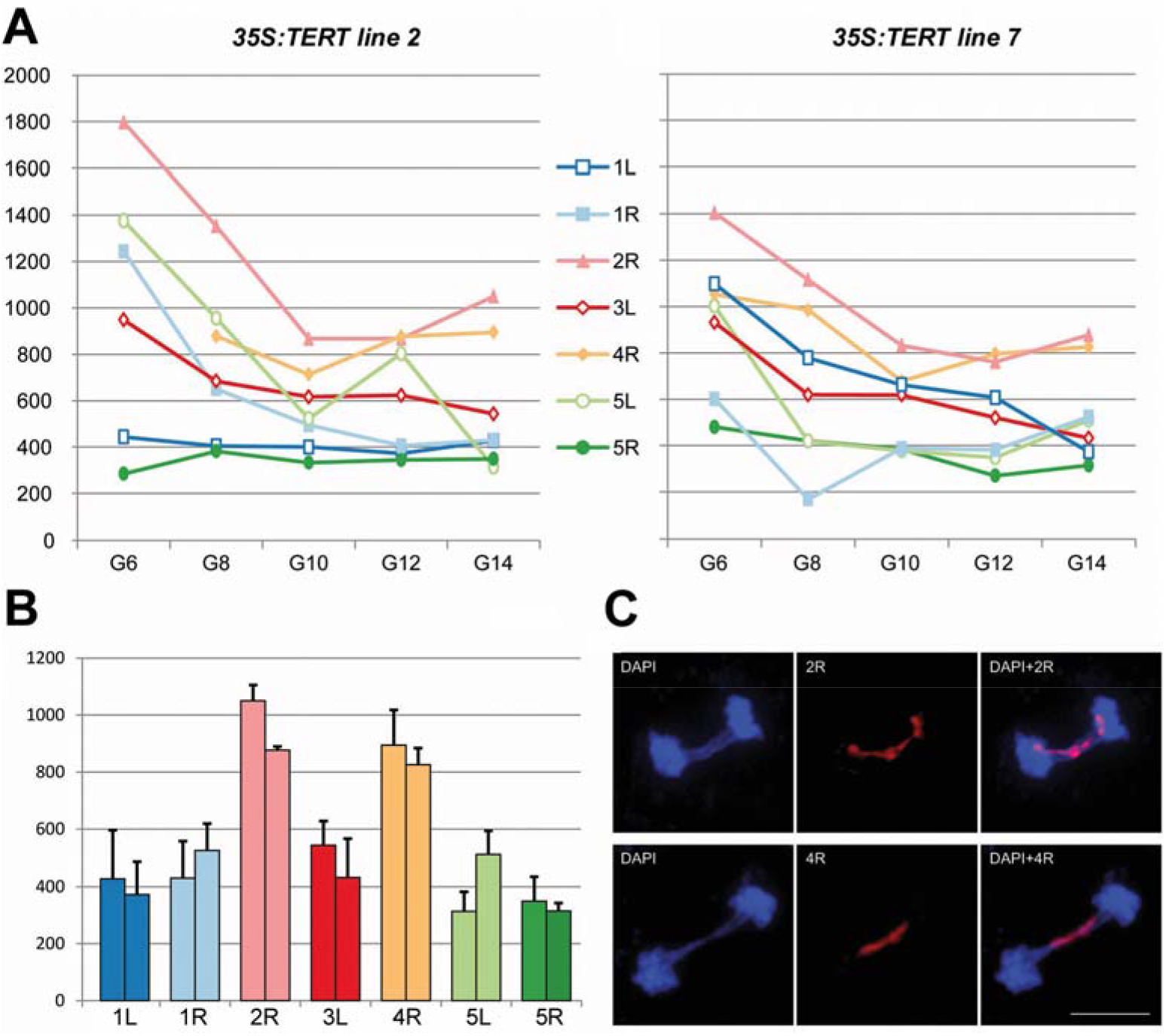
Analysis of individual telomeres in *35S:TERT* plants. (*A*) Measurement of individual telomere lengths over successive generations in lines 2 and 7 by PETRA. Measurements are the average of four plants, with the exception of G6/T2 in line where only one plant was available, and G12/T8 and G14/T10 where five plants were measured. (*B*) Comparison of telomere lengths in G14/T10 plants. Data is the same as in (*A*) but plotted as a bar graph for clarity. Error bars represent standard deviation for all data points in (*A*). (*C*) Representative figures of FISH analysis. Anaphase bridges showing telomere fusions of 2R and 4R are shown.

While the distribution of starting telomere lengths in the two lines varied from 300 to 1800 bp, the final distribution was strongly bimodal, with most telomeres clustered around 400 bp (Fig. 3B 1L, 1R, 3L, 5L, 5R; 421 ± 121 bp), and two telomeres clustered around 900 bp (Fig. 3B 2R, 4R; 910 ± 115 bp). Interestingly, despite varying starting lengths between the two lines, each individual telomere shortened to a similar final length in both lines. As apparent from Fig. 3A, the length of each telomere at G14/T10, was already reached in most cases by G10/T6, and for the shortest telomeres even from the beginning of the experiment in G6/T2. Although we terminated the experiment at generation G18/T14, the relative stability of telomeres from G10/T6 onwards implies that the hypomorphic telomerase was capable of maintaining telomeres at this length indefinitely. Our results further suggest that the hypomorphic telomerase is capable of maintaining telomeres only when they reach a chromosome-arm specific length. Since these late generation *35S:TERT* lines exhibit ongoing chromosome end-to-end fusions (Fig. 1D), it is likely that the ability of telomerase to extend particular telomeres directly correlates with their capping status; fully functional telomeres are not extendable, but telomeres that become partially deprotected are substrates for the hypomorphic telomerase. Along with the observation that further disruption of the telomerase pathway leads to lethality, we conclude that the *35S:TERT* lines maintain telomeres close to their minimal functional length.

The bimodal size distribution of telomere lengths in G14/T10 was intriguing, as it suggested that the length at which chromosome ends become accessible to the hypomorphic telomerase is telomere-specific. Therefore, we asked whether the two longer telomeres, 2R and 4R, were more stably capped than the others. To test this, we performed pairwise comparison of the frequency of anaphase bridges involving short and long telomeres by FISH in G14/T10 *35S:TERT* lines (Fig. 3C). As expected based on the phenotypic defects as well as the large numbers of anaphase bridges detected in late generation *35:TERT* plants, telomere to telomere fusions were readily detectable. Despite the large difference in telomere length, 2R and 4R were equally as likely to be involved in chromosome fusions as the shorter arms 1R and 1L (Table 1). This indicates that 2R and 4R telomeres become deprotected at longer lengths relative to the other chromosome arms.

**Table 1.** Dual label FISH analysis of anaphase bridges in Type II G14 /T10 mutants with probes recognizing either chromosome arms 4R and 5L, or 2R and 1R. No significant difference was observed between telomeres 4R (short) and 1L (long), and between 2R (long) and 1R (short).

|  | Anaphases with<br>bridges | Bridges with signal | Percentage | Fisher's exact<br>test |
| --- | --- | --- | --- | --- |
| 4R | 55 | 8 | 14,50 | 0,3879 |
| 1L | 55 | 6 | 10,9 |  |
| 2R | 52 | 20 | 38,5 | 0,34534 |
| 1R | 52 | 23 | 44,2 |  |

The dramatic difference in telomere length of 2R and 4R suggests that *cis* acting elements on these two subtelomeric arms regulate telomere homeostasis differently than on the other chromosome arms we have tested. We therefore examined telomere length in wild type plants to determine whether these two telomeres are differentially regulated under normal conditions as well. To accomplish this, we measured telomere length by STELA in 32 wild type plants from four bulk seed stocks. While in PETRA assay are the PCR adaptors anchored to the chromosome end via primer extension through telomeres, ligation is used in STELA (Capper et al., 2007, Heacock et al., 2004). We chose STELA over PETRA in this analysis due to concern that the primer extension reaction may not proceed through the entire length of wild type telomeres. Telomere length was measured as the length of the peak signal, and for allelic telomeres the length of the two alleles was averaged. As shown in Fig. 4A, the 2R and 4R telomeres were consistently longer than other telomeres in wild type plants as judged by two alternate methods. These data argue that *cis* acting elements affect the length of telomere homeostasis in the hypomorphic mutant as well as in wild type.

**Fig 4.**
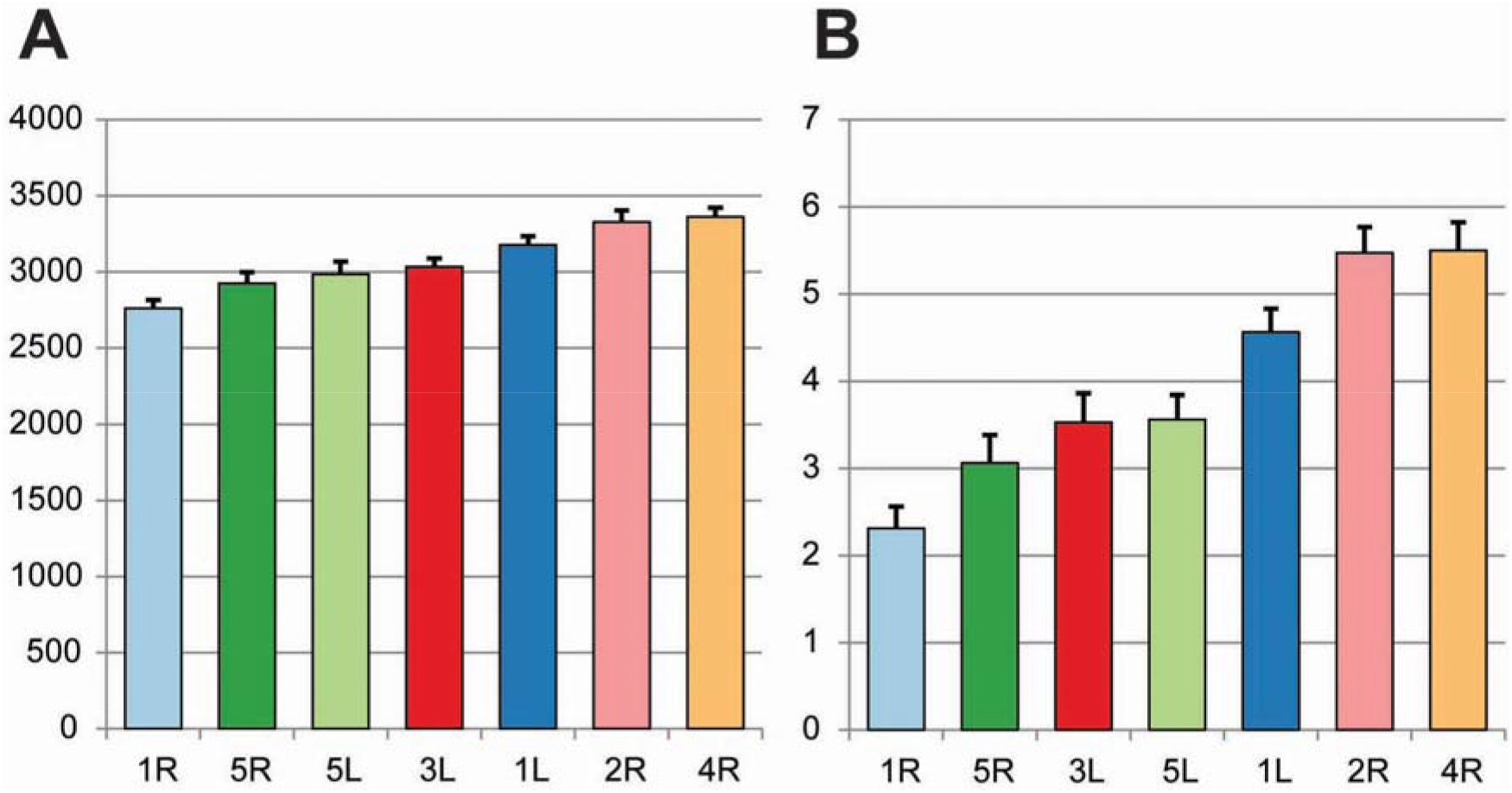
Single telomere analysis by STELA in wild type plants. (*A*) Telomere length as measured by STELA. Data represents the average of 32 plants. Error bars represent standard error of the mean. (*B*) Average telomere rank. As the average telomere length between plants varied, we ranked telomeres within each plant, telomeres were ranked from 1=shortest to 7= longest and the ranks were averaged. Error bars represent standard error of the mean.

## Discussion

In this study we describe a hypomorphic allele of TERT that produces a partially functional telomerase. Although this enzyme is incapable of maintaining telomeres within the wild type size range, it nevertheless sustains telomeres when they shorten to a length at which they become fusogenic. The enzyme also retains the capability to extend telomeres in plants depleted of Ku70. The Ku complex inhibits telomerase in Arabidopsis, and it also protects telomeres by preventing their exonucleolytic degradation and homologous recombination (Kazda, Zellinger et al., 2012, Riha & Shippen, 2003). Together, our data indicate that telomerase derived from the *35S:TERT* allele only accesses telomeres that are not fully capped. The molecular mechanism underlying the impaired telomerase performance in the *35S:TERT* lines is not known. It may be due to a reduced level of telomerase that is apparent from *in vitro* activity assay in comparison with wild type and *ACT2:TERT* plants. Notably, while the *35S:TERT* construct lacks introns, the *ACT2:TERT* construct contains an intron in the 5’UTR. It is well established that presence of introns enhances transgene expression in plants (Laxa, 2016), and therefore, the intronless *35S:TERT* construct may produce a suboptimal amount of TERT. Alternatively, the 35S promoter may not be efficiently active in tissues where telomerase is most needed, such as rapidly growing embryos (Sunilkumar, Mohr et al., 2002).

Although the hypomorphic *TERT* allele could not restore telomeres to wild type length, it was capable of maintaining telomeres at a shorter length set point. This is evidenced by the fact that *35S:TERT* plants can survive for 18 generations, and once the new shorter telomere length set point is established around G10/T6, plants maintain that telomere length for multiple generations. We confirmed that telomere length maintenance requires the catalytic activity of *TERT* since plants bearing a catalytically dead reverse transcriptase could not survive beyond the normal lifespan of *tert* mutants (G8-G10). Thus, while the hypomorphic *35S:TERT* allele is sufficient to indefinitely prolong plant lifespan, it can only sustain telomeres close to their minimal functional length. Additional support for this conclusion comes from the observation that in late generation *35S:TERT* lines deficient in the POT1A processivity factor lethality is immediate. We conclude that critically short telomeres in these lines cannot be maintained by telomerase and the plants succumb to telomere dysfunction.

The telomeres of late generation *35S:TERT* lines were maintained at a bimodal size distribution with arm-specific lengths corresponding to either 400 (average 421, arms 1L, 1R, 3L, 5L, and 5R) or 900 bp (910, arms 2R and 4R). FISH analysis unexpectedly demonstrated the shorter telomeres were no more prone to fusion than longer telomeres, implying that, despite their longer length, the capping status of the 2R and 4R telomeres is similar to other arms. Interestingly, the 2R and 4R telomeres are also maintained at longer lengths in wild type plants expressing normal TERT. Hence, cis-regulatory elements on these arms may not only modulate capping status, but also telomere length. A unique feature of the 2R and 4R telomeres is the immediate proximity of abundantly expressed protein coding genes (Vrbsky, Akimcheva et al., 2010), indicating that transcription of canonical transcripts may influence the status of neighboring telomeres. We note that regulation of telomere length by cis elements has been reported in humans (Britt-Compton, Rowson et al., 2006).

Despite extensive analysis of chromosome end protection in yeast and mammals, the relationship between the length of a telomere and its ability to serve as a functional cap on the DNA terminus remains enigmatic. Previous efforts to define the minimal functional length of telomeres have relied on identification of the shortest telomeres in a population of cells undergoing telomere crisis (Capper et al., 2007, Heacock et al., 2004, Heacock, Idol et al., 2007, Xu & Blackburn, 2007). A drawback to this approach is that cannot assess whether very short telomeres were captured as they were being processed by DNA repair machinery, or were the result of compromised checkpoint pathways. Other experiments have measured the amount of telomeric DNA retained at cloned chromosome fusion junctions, but in these cases it is impossible to determine telomere length prior to fusion (Heacock et al., 2004, Heacock et al., 2007). In contrast, in *35S:TERT* lines all telomeres are maintained at a length sufficient to confer long-term survival, but insufficient for buffering any additional loss of telomeric repeats.

For human cells, previous estimates of the minimal functional telomere size are between 42 (Xu & Blackburn, 2007) and 72 nt (Capper et al., 2007), lengths which are still sufficient for binding at least one TRF1 or TRF2 molecule. These observations, along with the fact that 72 nt of telomeric DNA bound by TRF2 could inhibit NHEJ *in vitro* (Bae & Baumann, 2007), have led to the proposal that the minimal functional length is defined by the ability to bind TRF2. Similarly, yeast cells continue to grow and divide until telomeres reach less than 100 bp (Forstemann et al., 2000). Our results argue that the minimal functional length in Arabidopsis is significantly longer at approximately 400 bp, consistent with previous analysis of telomere fusion junctions reporting a minimal functional length of 360 bp (Heacock et al., 2004). While several telomere-specific binding components have been described for Arabidopsis (Schrumpfova, Vychodilova et al., 2014, Song, Leehy et al., 2008, Surovtseva, Churikov et al., 2009), the limited biochemical data available for these factors suggest the binding sites are similar to the yeast and mammalian telomere proteins, and hence much smaller than 400bp. In contrast, 400 bp is the approximate length required for circularization of double-stranded DNA. Experiments on naked DNA have suggested that DNA molecules with lengths of roughly 300-400 bp are optimal for circular ligation (Shore, Langowski et al., 1981). In addition, *in vitro* experiments with TRF2 revealed that telomeric tracts of 500 bp are sufficient to form t-loops (Griffith et al., 1999, Stansel, de Lange et al., 2001). Although the length of looped DNA in a t-loop is proportional to total telomere length, the loops in the 500 bp artificial constructs averaged 338 bp, very close to the minimal telomere length detected at multiple chromosome arms in this study. Therefore, we postulate that formation of a secondary structure such as a t-loop is required for proper capping of chromosome ends, and thus defines the minimal functional length for telomeres in Arabidopsis.

## Materials and Methods

### Plants and Growth Conditions

Plants were grown in soil under long day conditions with 16 hr daylight at 21°C. All mutants and genotyping conditions have been previously described: *tert* (Fitzgerald, Riha et al., 1999), *ku70* (Riha et al., 2002), *pot1a* (Surovtseva et al., 2007).

### Constructs and plant transformation

Full length *TERT* cDNA was obtained from reverse-transcription reactions from wild type RNA using the RevertAid First Strand cDNA Synthesis Kit and amplified with Phusion Hot Start II polymerase (Fermentas) using primers TERT cDNA 5’ BamHI (5’-GGATCCAAGGAGGAGGAAGGTGTAATG-3’) and TERT cDNA 3’ SacI (5’-GAGCTCAAATGTTACAAATCCATTTTG-3’). Full length cDNA products were subcloned into pCR2.1-TOPO (Invitrogen) and sequenced. The cDNA was then transferred to binary vectors pCBK05 (35S promoter) forming pCBJ05 or pCBM10 (Actin promoter) forming pCBJ06 using BamHI and SacI sites. For the D680N mutation, pCBJ05 was digested with HindIII, releasing a small fragment containing the catalytic residue, and subcloned into pBluescript. Mutagenesis was performed using the QuikChange site directed mutagenesis kit (Stratagene) using manufacturer’s instructions and the primers AtD860NF(5’-GAGATTTATTAATGACTACCTTTTTG-3’) and AtD860NR (5’CAAAAAGGTAGTCATTAATAAATCTC-3’) resulting in a change from GAT to AAT. The mutated fragment was re-inserted into pCBJ05. Transformation of G4 *tert* mutants was performed by the floral dip method and transformed plants were selected for resistance to the herbicide Basta.

### TRAP assay

TRAP assays were performed as previously described (Kannan, Nelson et al., 2008).

### Telomere length analysis

DNA extraction and TRF analysis were performed as previously described (Watson & Shippen, 2007). For both PETRA (Heacock et al., 2004) and STELA (Baird, Rowson et al., 2003) 20 ng of total genomic DNA were used. For PETRA, the PETRA-T primer (5’-CTCTAGACTGTGAGACTTGGACTACCCTAAACCCT-3’) was annealed to G-overhangs and extended in a 20 μL reaction containing 1x Phi29 buffer (Fermentas), 125 μM dNTPs, 0.5 μM PETRA-T, and 4 U Phi29 at 30°C for 1 hr. Samples were heat inactivated at 65°C for 20 minutes, precipitated with ethanol, and resuspended in 20 μL of water. For STELA, the primer Fok2 (5’-CTCTAGACTGTGAGACTTGGACTACAGGATGTAAACCC-3’) was ligated to genomic DNA in 20 μL reactions containing 1x T4 DNA ligase buffer, 0.5 μM Fok2, and 1 μL T4 DNA ligase (Fermentas) overnight at 16°C. Samples were heat inactivated at 65°C for 20 min, precipitated with ethanol, and resuspended in 20 μL of water. Following this initial processing, STELA and PETRA reactions were identical. 1 μL of STELA or PETRA products were PCR amplified in 20 μL reactions containing 1x Phusion GC buffer, 200 μM dNTPs, 0.5 μM PETRA-A (5’-CTCTAGACTGTGAGACTTGGACTA C-3’), 0.5 μM of one subtelomere specific primer (1L: 5’-AGGACCATCCCATATCATTGAGAGA-3’, 1R: 5’-CTATTGCCA GAACCTTGATATTCAT-3’, 2R: 5’-CAACATGGCCCATTTAAGATTGAACGGC-3’, 3L: 5’-CATAATTCTCACAG CAGCACCGTAGA-3’, 4R: 5’-TGGGTGATTGTCATGCTACATGGTA-3’, 5L: 5’-AGGTAGAGTGAACCTAACA CTTGG-3’, 5R: 5’-CAGGACGTGTGAAACAGAAACTACA-3’), 0.4 U Phusion Hot Start II (Fermentas). PCR reactions were performed by incubating samples at 98°C for 3 min followed by 22-25 cycles of 98°C 15 sec, 60°C 30 sec, 72°C 2 min 30 sec, followed by a final incubation at 72°C for 10 min. Reactions were then separated on 1.2% agarose gels cast in 12×14 cm trays at 100V for 2 h 30 min. DNA was transferred to nylon membranes by Southern blotting. Hybridization was performed as previously described (Heacock et al., 2004), membranes were exposed to Kodak phosphoscreens, visualized with a Pharos FX Plus (Biorad) and images were analyzed with Imagelab (Biorad).

### Chromosome Preparation, Probe Labeling, BAC FISH and Image Processing

Whole inflorescences of *Arabidopsis thaliana* were fixed in freshly prepared ethanol:acetic acid fixative (3:1) overnight, transferred into 70% ethanol and stored at −20°C until use. Selected inflorescences were rinsed in distilled water and citrate buffer (10 mM sodium citrate, pH 4.8), and digested by a 0.3% mix of pectolytic enzymes (cellulase, cytohelicase, pectolyase; all from Sigma) in citrate buffer for c. 3 hrs. Chromosome preparations were prepared from pistils and pretreated with RNase (100μg/ml, AppliChem) and pepsin (0.1mg/ml, Sigma-Aldrich) as previously described (Mandakova, Marhold et al., 2014). *Arabidopsis thaliana* BAC clones T25K16 (AC007323), F6F3 (AC023628), F22L4 (AC061957), T1N6 (AC009273) and F22N6 (B98711) were used for *in situ* identification of chromosome arm 1L, F19K16 (AC011717), F18B13 (AC009322), F5I6 (AC018848), T21F11 (AC018849) and F23A5 (AC011713) for 1R, F13A10 (AC006418), T3A4 (AC005819), F19D11 (AC005310), F14M4 (AC004411) and T8I13 (AC002337) for 2R, and F19H22 (AL035679), T22F8 (AL050351), F23K16 (AL078620), T23P19 (B98327) and T5J17 (AL035708) for 4R. DNA probes were labeled with biotin-dUTP and digoxigenin-dUTP by nick translation as previously described (Mandakova, Joly et al., 2010). Differentially labeled BACs corresponding to 1L+4R and 1R+2R chromosome arms were pooled, precipitated, and resuspended in 20μl of hybridization mixture (50% formamide and 10% dextran sulfate in 2×SSC) per slide. Probes and chromosomes were denatured together on a hot plate at 80°C for 2 min and incubated in a moist chamber at 37°C overnight. Post hybridization washing was performed in 20% formamide in 2×SSC at 42°C. Fluorescent detection was as follows: biotin-dUTP was detected by avidin–Texas Red (Vector Laboratories) and amplified by goat anti-avidin–biotin (Vector Laboratories) and avidin–Texas Red; digoxigenin-dUTP was detected by mouse antidigoxigenin (Jackson ImmunoResearch) and goat antimouse Alexa Fluor 488 (Molecular Probes). Chromosomes were counterstained with DAPI (4’,6-diamidino-2-phenylindole; 2 μg/ml) in Vectashield (Vector Laboratories). Fluorescence signals were analyzed with an Olympus BX-61 epifluorescence microscope and CoolCube CCD camera (MetaSystems). Images were acquired separately for the two fluorochromes using appropriate excitation and emission filters (AHF Analysentechnik). The two monochromatic images were pseudo colored and merged using Adobe Photoshop CS2 software (Adobe Systems).

## Acknowledgements

The work was supported from the European Regional Development Fund-Project „REMAP” (No. CZ.02.1.01/0.0/0.0/15_003/0000479), by the Czech Science Foundation (18-20134S), by EMBO Installation Grant (1304130933), by the Austrian Academy of Sciences and by the National Institutes of Health (GM R01-GM065383).

