## Supplementary Figures for "A hypomorphic allele of telomerase reverse transcriptase uncovers the minimal functional length of telomeres in Arabidopsis"

**Supplementary data**

**Fig S1.**

Schematic showing the generation of the *35S:TERT* and *35S:TERTDN680N* lines used in this study. *Actin:TERT* lines were generated in the same fashion as the *35S:TERT* lines.

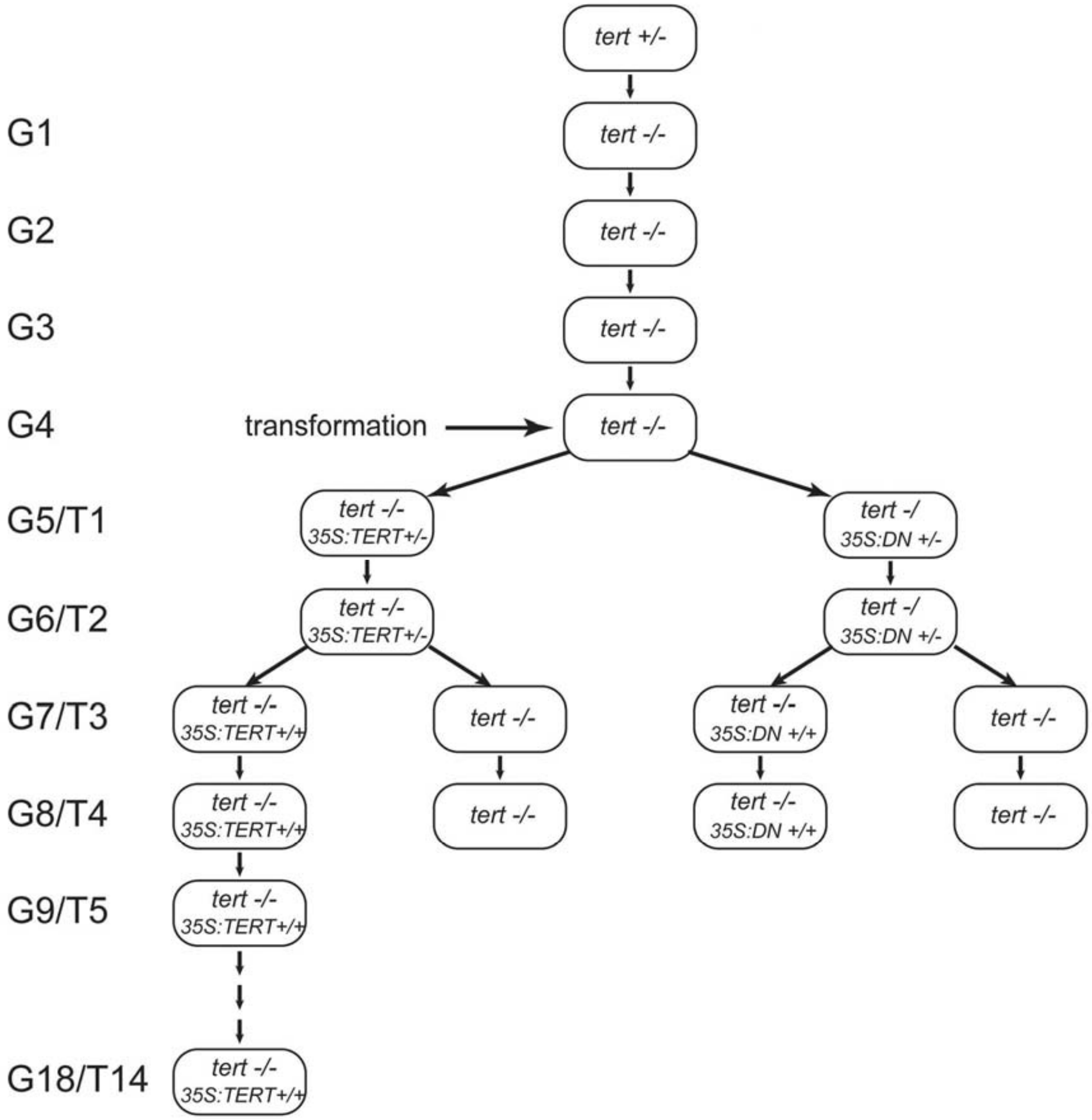

**Fig S2**

Populations of multiple generations of *35S:TERT* line 7 plants, from G10/T6 through G16/T12. No obvious growth or developmental difference is observed with progressive generations at the population level.

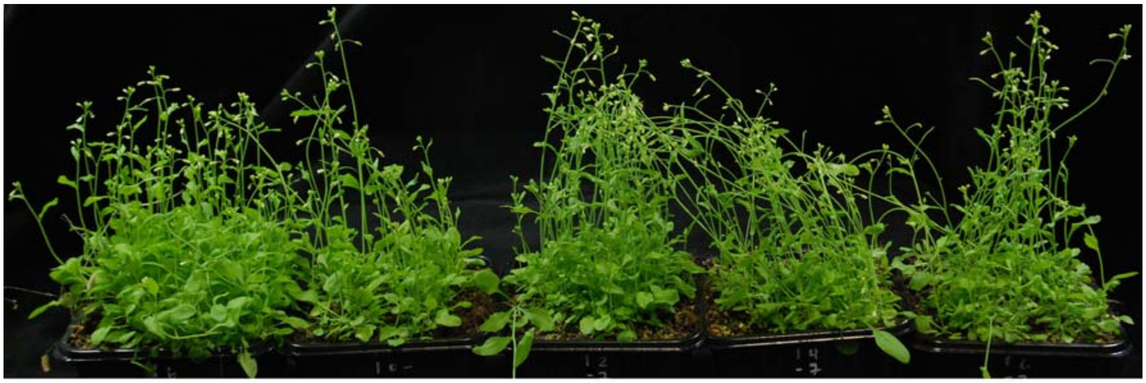

wild type G10/T6 G12/T8 G14/T10 G16/T12

**Fig S3**

TRAP assay results for the *35S:TERT(D680N)* dominant negative mutant.

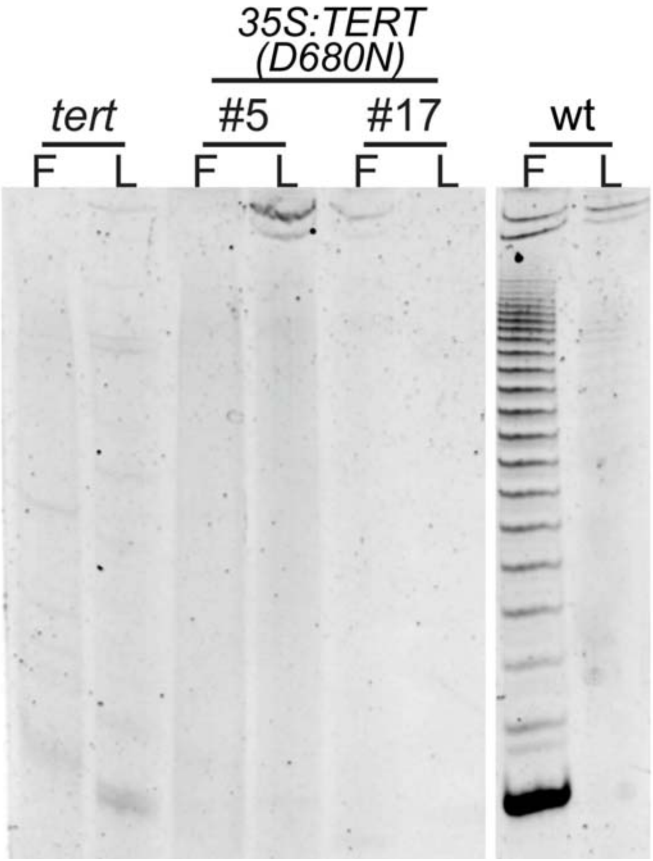

**Fig S4**

Schematic displaying the generation of *tert ku70 35S:TERT* lines.

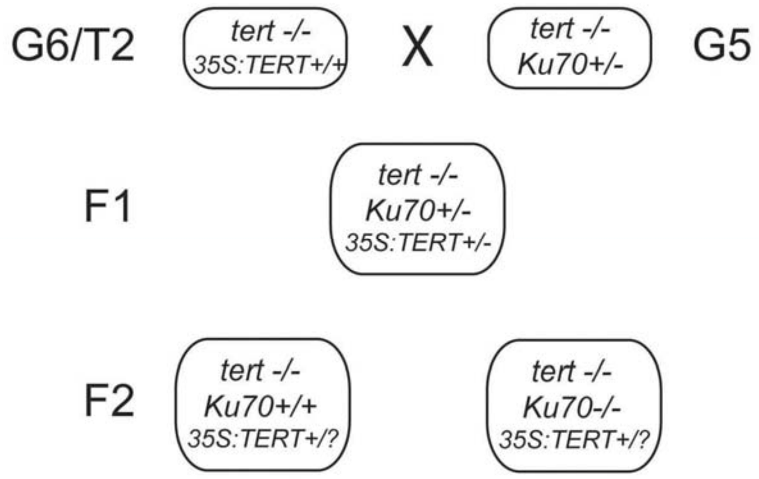

**Fig S5**

Schematic displaying the generation of the *tert pot1a 35S:TERT* lines.

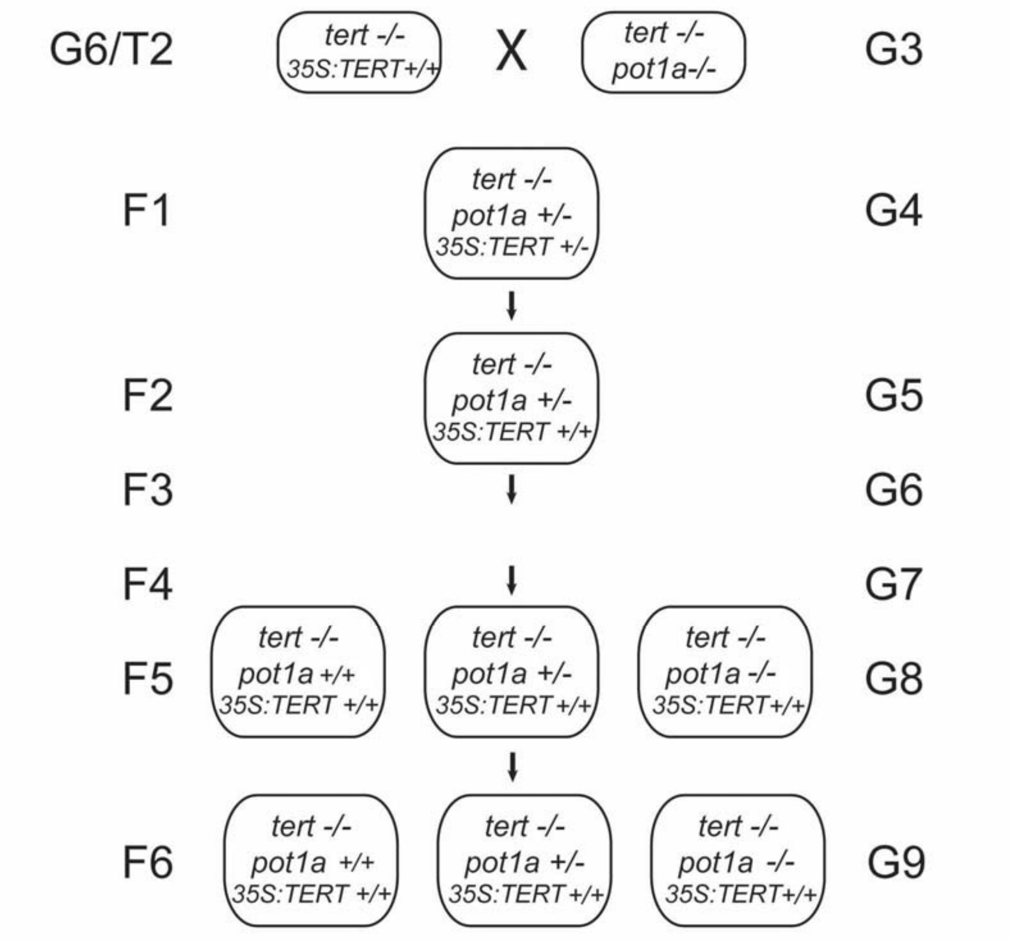
